# Metabolomics reveals synergistic antimalarial drug pairing effects against *Plasmodium falciparum in vitro*

**DOI:** 10.64898/2025.12.13.694081

**Authors:** Francois Eya’ane Meva, Tarrick Qahash, Justin Arinaga, Manuel Llinás

## Abstract

**Background:** The development of new drugs against afflictions that disproportionately impact poorly resourced areas around the globe is an expensive endeavor. As cost-effective alternatives, strategic combinations of approved drugs can be used to enhance the efficacy against *Plasmodium falciparum*. Understanding the metabolic consequences of such combinations is essential for optimizing treatment strategies and delaying drug resistance.

**Methods:** An integrated metabolomic and pharmacological analysis was performed on P. *falciparum* exposed to chloroquine (CQ), pyrimethamine (PY), sulfadoxine (SD), and their combinations (SDPY and SDCQ). Dose□response assays were used to quantify drug potency, whereas untargeted metabolomic profiling was used to assess pathway-level perturbations associated with individual and combined treatments.

**Results:** Dose**□**response assays confirmed the nanomolar potency of PY (IC₅₀ = 12.5 nM) and CQ (IC₅₀ = 11 nM) compared with the micromolar efficacy of SD (IC₅₀ = 9.1 µM), which is consistent with its role as a synergistic antifolate partner. Metabolomic profiling revealed that PY strongly disrupted folate-dependent pyrimidine biosynthesis, leading to deoxyuridine and dUMP accumulation, whereas SD caused milder perturbations, which was consistent with DHPS inhibition. CQ produced modest metabolic effects alone but markedly amplified antifolate-induced stress when combined with PY.

Drug combinations generated metabolic responses that are distinct from those resulting from individual treatments. Across antifolate combinations, consistent trends included reduced amino acid pools, suppression of thiamine and glutathione metabolism, and enhanced PPP inhibition, leading to broad disruption of nucleotide, redox, and carbon metabolism.

**Conclusions:** Pyrimidine suppression has emerged as the central hallmark of antifolate-based therapy in *P. falciparum*. The distinct synergistic signatures observed with drug combinations support their potential to enhance efficacy and delay resistance. These findings provide a mechanistic foundation for guiding antimalarial combination policies, optimizing therapeutic regimens, and strengthening rational drug-design strategies.

## Introduction

Malaria is an acute febrile illness caused by *Plasmodium* parasites, which are spread to people through the bites of infected female *Anopheles* mosquitoes. The eukaryotic protozoan *Plasmodium* parasites have a complex life cycle that imposes a major health and economic burden in endemic countries [1]. In 2024, there were more than 280 million estimated cases of malaria and more than 600,000 malaria-related deaths across 80 malaria endemic countries with sub-Saharan Africa accounting for the majority of new infections [1]. Alongside preventive measures, chemotherapy continues to be a critical component of malaria control, complementing vector control strategies and efforts to improve access to effective treatment [2]. The WHO advises the use of malaria vaccines to protect children in malaria-endemic regions, with priority given to settings experiencing moderate to high transmission. The two vaccines currently recommended are RTS,S and R21/Matrix-M (R21)[1]. On a positive note, the WHO’s Strategic Advisory Group of Experts on Immunization (SAGE) and the Malaria Policy Advisory Group (MPAG) favorably reviewed the RTS,S/AS01 malaria vaccine study, which showed that a four-dose schedule reduced severe malaria by 54%, with the fourth dose adding a 30% benefit over three doses [1, 3]. Unfortunately, many African countries continue to face high transmission and mortality rates, compounded by the growing threat of antimalarial drug resistance and the declining efficacy of artemisinin-based therapies [1, 4]. Concurrently, genetic changes in parasites and increasing insecticide resistance are undermining diagnostic reliability and vector control efforts, which threatens to reverse years of progress [1].

Treatment of malaria has faced extreme challenges with the emergence and spread of drug-resistant *P. falciparum* strains for most commercially available antimalarial drugs, creating a heavy burden on public health and economic development [5]. A major strategy for overcoming clinical parasite resistance is based on the use of drug combinations [6]. However, the modes of action of many drug combinations remain understudied.

Chloroquine (CQ) was first used as a front-line drug by sub-Saharan African countries in the early 20^th^ century [7, 8]. Sulfadoxine (SD) and pyrimethamine (PY) are antifolates used to treat uncomplicated malaria [9]. CQ was later replaced by sulfadoxine-pyrimethamine (SDPY) due to parasite resistance to CQ monotherapy [7]. CQ binds tightly to ferriprotoporphyrin IX (FPIX) to form the toxic FPIX-CQ complex, which inhibits the formation of hemozoin, poisons the acidic digestive vacuole (DV), and starves the parasite [10]. The catabolism of hemoglobin in the DV provides critical amino acids to the parasite, releasing free heme, whereas the folate biosynthesis pathway delivers the building blocks for DNA synthesis [11].

*P. falciparum* resistance to CQ is associated with mutations in the pfcr gene, which encodes the chloroquine resistance transporter (PfCRT) [12]. SD and PY are inhibitors of dihydropteroate synthase (DHPS) in the folate biosynthetic pathway. DHPS couples p-aminobenzoic acid (PABA) to 7,8-dihydropterin, producing dihydropteroate in the step preceding dihydrofolate synthesis [13]. Previous studies have identified mutations at four codons (N51, C59, S108, and I164) that confer resistance to PY, whereas five DHPS codons (S436A/F; A437G; K540E; A581G; and A613S/T) are associated with SD resistance in *P. falciparum* [14, 15]. An effective strategy for reducing parasite resistance is to use a combination of drugs for synergistic *effects* or 1+1 > 2 *effects* [16]. Hence, therapy with SDPY and CQ in Ugandan infants and young children has been more effective than monotherapy in clearing parasites and improving cure rates (88% combined vs. 55% for chloroquine alone) [17].

The efficacy of combining drugs can also be studied via isobolograms [15]. Isobolograms from combined treatment with 4-aminoquinoline and dihydroartemisinin derivatives revealed substantial synergistic interactions between these drugs. In this case, the potential to inhibit hemozoin formation is enhanced by the combination, compared with the individual compounds evaluated individually. The reduction in hemozoin due to 4-aminoquinolines is more pronounced in combination with dihydroartemisinin [18]. Isobologram studies have also demonstrated that the synergy between PY and SD depends strongly on folate levels, with PY acting beyond DHFR inhibition, potentially blocking folate uptake or use, and engaging additional targets that restore DHPS as a relevant site for SD, thereby markedly suppressing parasite growth [19].

Metabolomics is a comprehensive approach to simultaneously capture a broad array of metabolites that has transformed the study of the metabolism of infectious disease agents [20, 21]. The application of metabolomics in the malaria field has enabled the characterization of both established and potential future antimalarials, thus contributing to rational antimalarial drug design [22]. Metabolite profiling via mass spectrometry is increasingly being used to measure the metabolic perturbations induced by antimalarial compounds [23, 24, 25]. In the case of *P. falciparum*-induced malaria, the effects of widespread erythrocyte lysis and ischemic damage arising from the sequestration of parasitized erythrocytes within the microvasculature coupled with the high metabolic demands of the rapidly proliferating parasite are largely responsible for the pathogenesis of the disease and its manifestations [26].

The advantages of using paired drug combinations have long been realized and are based on two major principles. First, the starting frequency of malaria parasites resistant to both drugs in a population is significantly reduced. Second, multidrug-resistant alleles are lost more frequently during recombination during meiosis [27]. In this work, we have characterized the metabolic effect of treating *P. falciparum* parasites with paired combinations of sulfadoxine, pyrimethamine, and chloroquine to identify efficacious drug synergies.

## Materials and methods

### Antiplasmodial assay

*P. falciparum* blood-stage parasites of strain 3D7 were cultured according to standard protocols at 37°C, 5% CO_2_, 5% O_2_, 90% N_2_ and 80% humidity in complete RPMI medium with 0.45% (w/v) AlbuMAXII, 0.2 mM hypoxanthine, 25 μg/mL gentamicin and human A erythrocytes as previously described [28]. The volume of standard cultures was 14 mL, and the hematocrit was 3% unless otherwise stated. Synchronized 3D7 parasites were obtained after treatment with 5% sorbitol [29]. IC_50_ values were determined using a SYBR Green 1 assay according to established protocols [30, 31]. Media of 1% parasitemia, 1% hematocrit cultures and the drug were incubated for 72 h in 96-well plates. The plates were then wrapped with parafilm and stored overnight at −80°C. The plates were thawed at room temperature, and an aliquot (100 µL) of buffered SYBR Green (Molecular Probes, Inc., Eugene, OR) was added to each culture-containing well using a multichannel Pipetteman. Mixing was achieved by pipetting up and down until no cell sediment remained. The plate was wrapped in aluminum foil and stored in an incubator for 6 h. DNA quantification was performed using a TecanGENios microplate detection device.

#### Sample preparation for antimalarial metabolite profiling

Chloroquine diphosphate (chloroquine) 98% was purchased from Fisher Scientific. Pyrimethamine 98% and sulfadoxine 98% were obtained from Sigma□Aldrich. All drugs were stored as dry solids at room temperature and prepared as concentrated stock solutions in DMSO. Drug treatments were performed according to Allman *et al*.(2016), with some modifications. Atovaquone treatment was used as a positive control, and an untreated control was included. Briefly, magnetically purified parasites (20 to 30 μL pellets) were incubated in 6-well plates at 0.5% hematocrit with antimalarial compounds at 10 × IC_50_ for 2.5 h [32].

#### Metabolite extraction

Extractions were performed as described previously [33, 34]. Briefly, 20 μL pellets enriched with parasitized cells were resuspended in 1.0 ml of prechilled 90:10 methanol:water and placed at 4°C. The internal standard [^13^C_4_,^15^N_1_] aspartate was spiked into the extraction methanol solution to control for sample preparation and handling. The samples were vortexed, resuspended, and centrifuged for 10 min at 15000 rpm and 4°C. The dried metabolites were resuspended in HPLC-grade water (Chromasolv; Sigma) to a concentration between 1.0 × 10^5^ and 1.0 × 10^6^ cells/μLon the basis of hemocytometer counts of purified parasites. All samples were processed in triplicate with method blanks to reduce technical variation and account for background signals [32].

#### LC**□**MS instrumentation

Liquid chromatography**□**mass spectrometry (LC**□**MS) was performed using anExactive Plus system and a XSelect HSS T3 column (100 Å, 2.5 µm, 2.1 mm X 100 mm). In all instances, the data presented are the means ± standard deviations for N = 3 biological replicates. Data extraction, treatment, periodicity and microarray correlation analyses are described in the Supplemental Methods [35].

#### Metabolomics data analysis

Raw data files from the ThermoExactive Plus orbitrap (.raw) were converted to a format compatible with our analysis software (.raw/.mzXML) [36, 37]. Spectral data (.mzXML files) were visualized in MAVEN, a freely available software package for MS-based metabolomics [38]. Data were peak picked using a permissive threshold (S/N = 3) and identified on the basis of m/z within 15 ppm and retention times consistent with a curated standard library. The extracted ion chromatogram of each signal was visually inspected, and data originating from peak picking errors, thermal noise, elution artifacts, or those associated with the void and wash volumes were excluded. Coeluting adducts, fragments, and isotopomers were condensed into their respective parent masses, and the intensities for each of the final parent masses were hand verified to correct for peak picking errors [39]. The metabolites were examined using their log_₂_ fold changes and a two-dimensional hexagonal metaprint by arranging the most influential nodes on the outer edge of the suprahexagon to enable comparative profiling [32].

## Results

**The** IC_50_ values for PY, CQ, and SD were determined for synchronous cultures of the chloroquine-sensitive *Plasmodium falciparum* strain 3D7 (Figure 1). The results confirmed that parasites appeared to be more sensitive to CQ, suggesting compound-specific metabolic dependencies. These findings are in line with previous reports showing divergent activity profiles of antifolate and quinoline drugs in *P. falciparum* [40]. On the basis of these experiments, IC_50_ values of 9.1 µM for SD, 11 nM for CQ, and 12.5 nM for PY were determined for profiling.

**Figure 1.**
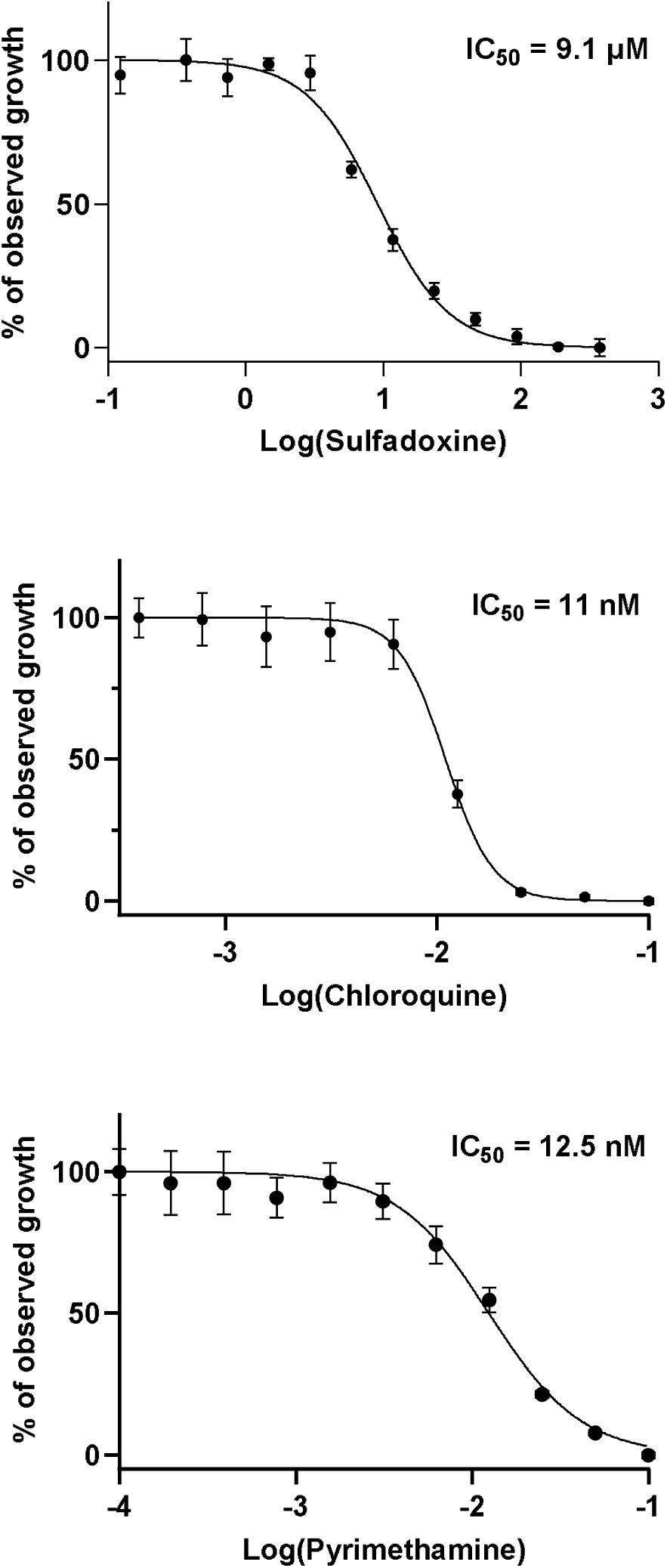
Dose□response curves of *Plasmodium falciparum* exposed to antifolate and quinoline drugs. Parasite growth inhibition was measured following treatment with increasing concentrations of PY, CQ, and SD. The data are expressed as the percentage of observed growth relative to that of untreated controls. Nonlinear regression analysis was performed to estimate the IC₅₀. PY displayed an IC₅₀ of 12.5 nM, CQ had an IC₅₀ of 11 nM, SD had an IC₅₀ of 9.1 μM. The error bars represent the means ± sems of triplicate assays.

### Global metabolic profiling by liquid chromatography–mass spectrometry (LC□MS)

Metabolic changes were profiled using LC-MS following drug exposure at 10 x IC50 concentrations for drugs alone and for the paired drug combinations of sulfadoxine (SD)-chloroquine (CQ) and sulfadoxine (SD)-pyrimethamine (PY). Significant alterations were observed across several metabolic pathways, including perturbations of nucleotides (Figure 2), pyrimidine biosynthesis (Figure 3), the central carbon/TCA cycle and energy metabolites (Figure 4), amino acid metabolism (Figure 5), coenzyme and vitamin metabolism (Figure 6), hemoglobin digestion (Figure 7),and the pentose phosphate pathway (Figure 8). A global heatmap of 154 metabolites is depicted in Figure 9.

**Figure 2.**
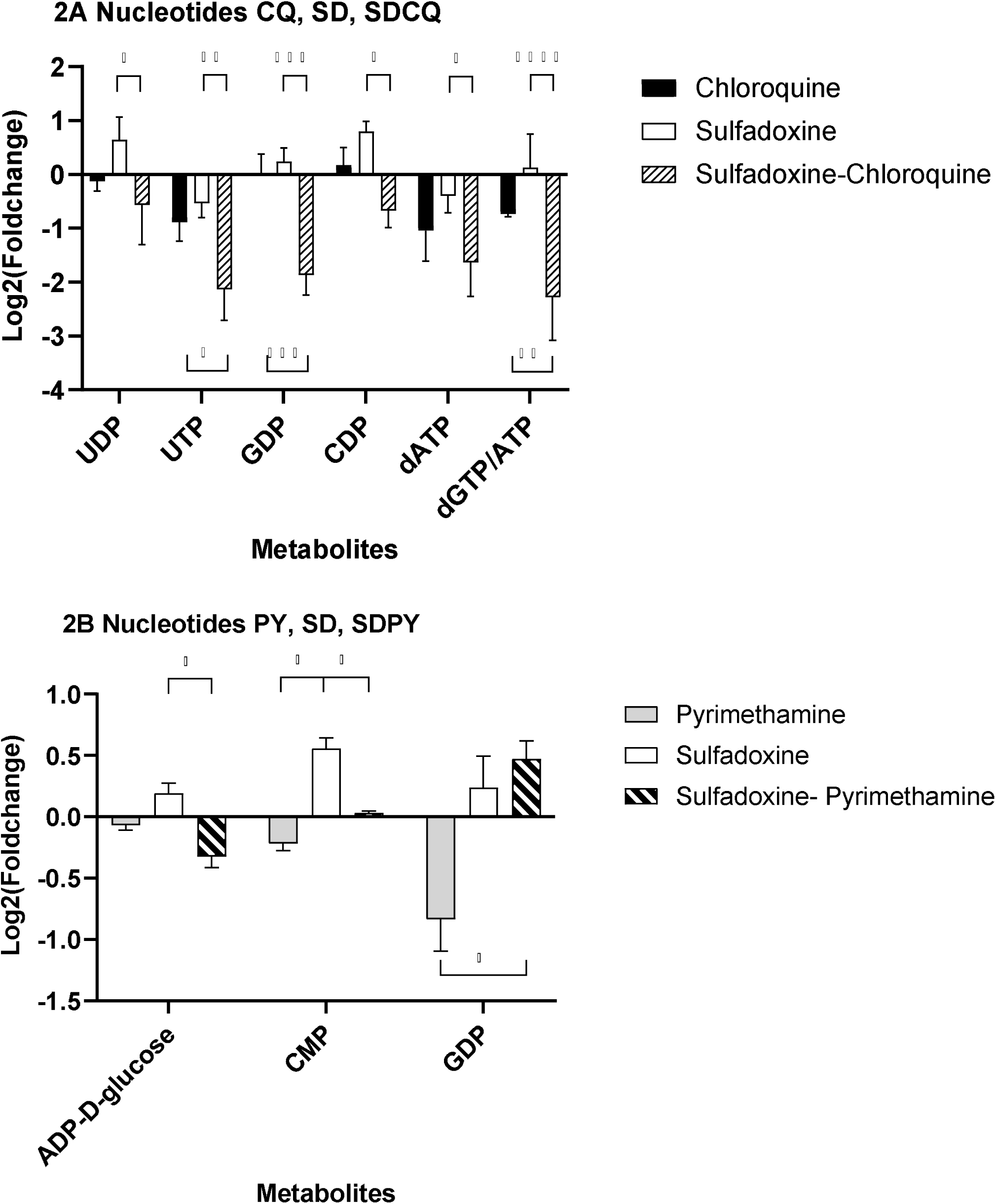
Nucleotide metabolites from parasite treatment with individual drugs and their combinations SDCQ, and SDPY. A) SDCQ: CQ alone produced marked dCDP depletion. SD alone induced notable increases in CMP and ADP-D-glucose, consistent with the accumulation of pyrimidine-related intermediates. Compared with SD alone, the combination of SDCQ neutralized these accumulations, with CMP returning near baseline and ADP-D-glucose decreasing. B) SDPY: PY alone reduced GDP and ADP-D-glucose. SD alone caused increases in CMP and ADP-D-glucose. Compared with PY alone, the SDPY combination resulted in more pronounced depletion of ADP-D-glucose, normalization of CMP levels, and an increase in GDP.

**Figure 3.**
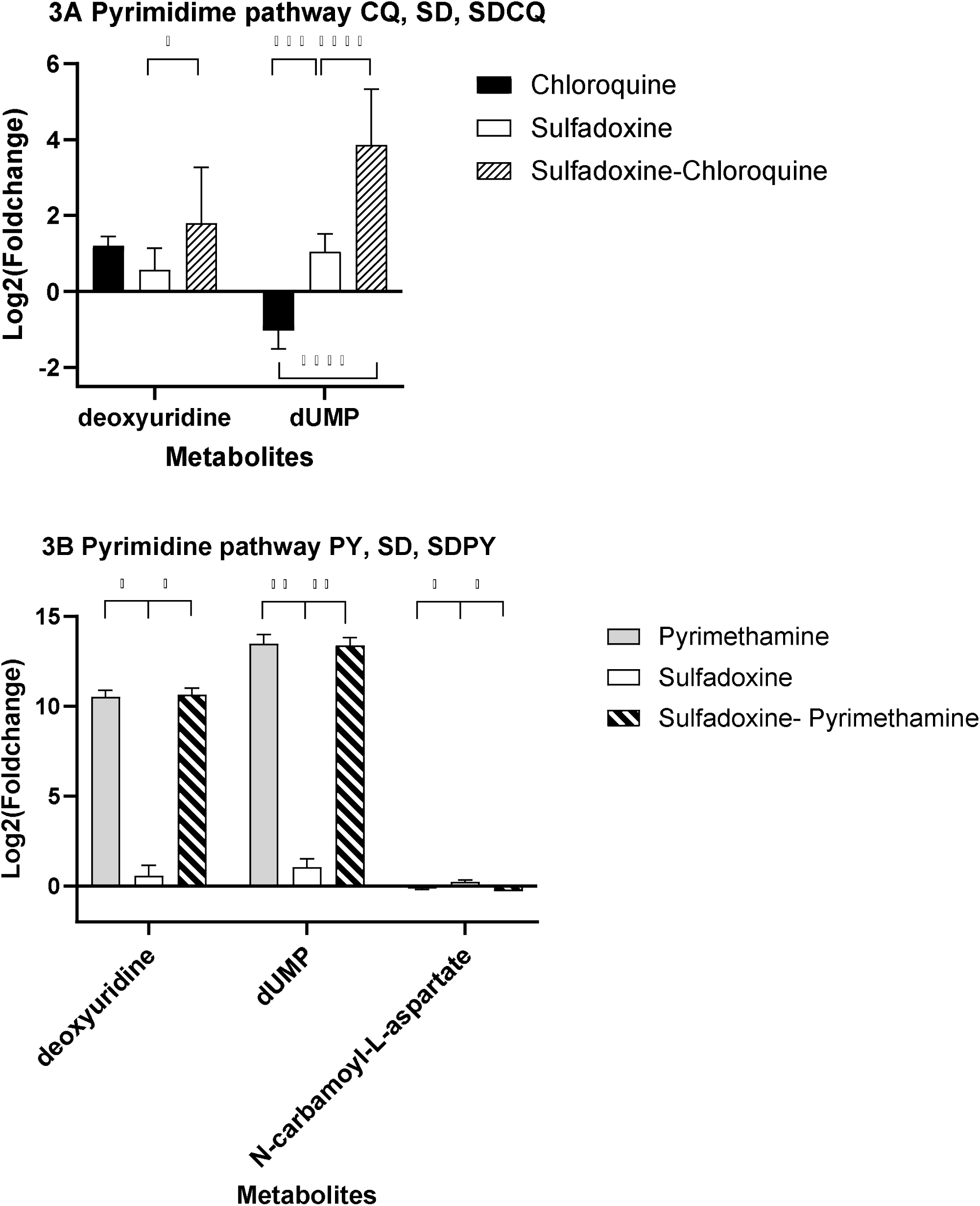
Pyrimidine metabolites in the case of isolated drugs and their combinations SDCQ and SDPY. A) SDCQ: CQ alone yielded mild increases in deoxyuridine and depletion of dUMP. SD alone induced a mild elevation of deoxyuridine and caused modest dUMP accumulation. SDCQ combination resulted in the accumulation of deoxyuridine and marked recovery of dUMP levels (1.5–6.5-fold). B) SDPY: PY alone caused extreme accumulation of deoxyuridine (≈10-fold) and dUMP (13-14-fold). SD alone moderately elevated deoxyuridine, resulting in modest dUMP increases (0.5–2-fold) and near baseline accumulation of N-carbamoyl-L-aspartate. The SDPY combination reproduced the high levels of deoxyuridine and dUMP typical of PY while suppressing N-carbamoyl-L-aspartate below baseline.

**Figure 4.**
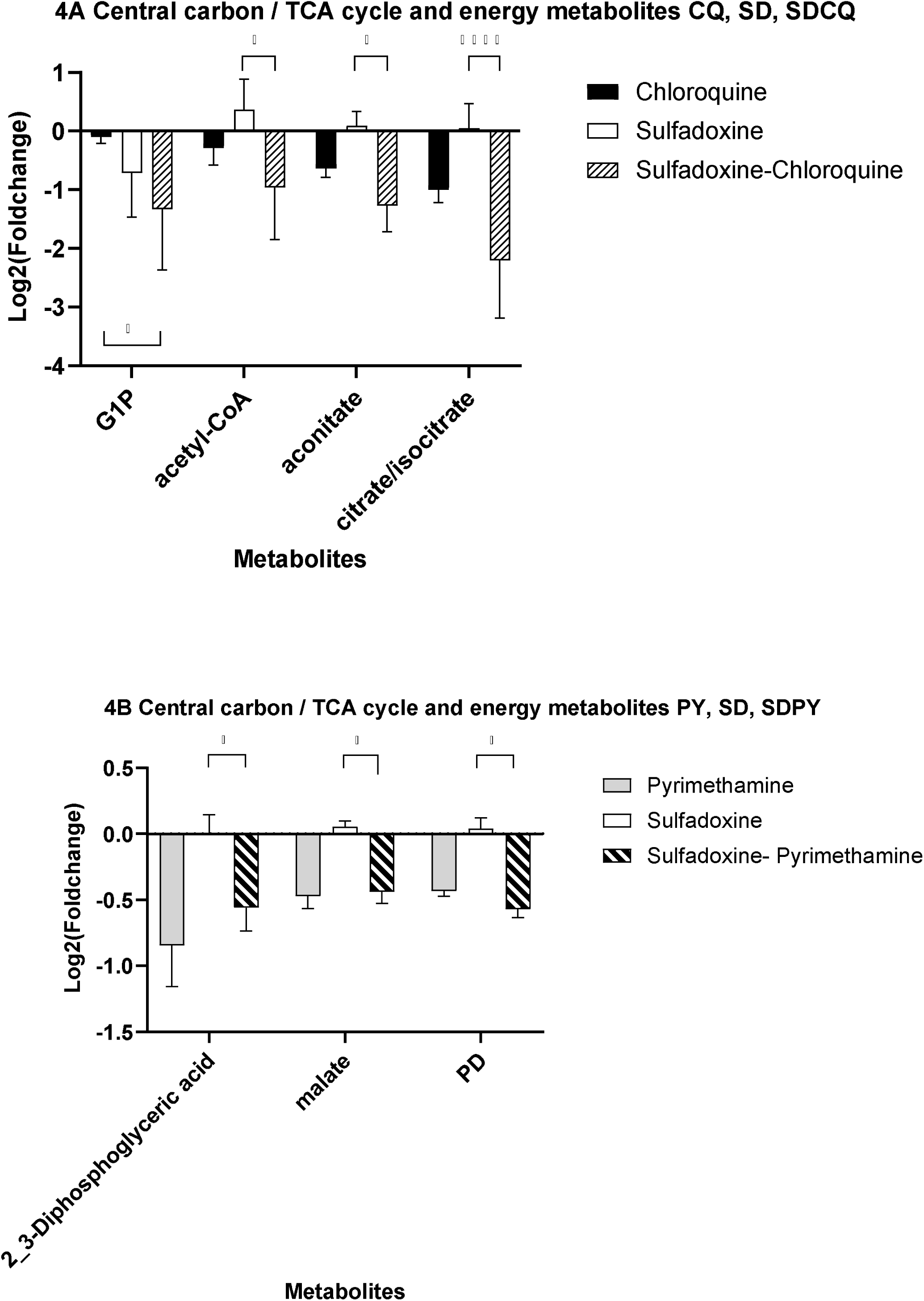
Central carbon/TCA cycle and energy metabolites in the case of isolated drugs and their combinations SDCQ and SDPY. A) SDCQ: CQ alone caused broad depletion of glycolytic/TCA intermediates: G1P (0.08 to −0.11), acetyl-CoA (0.29 to −0.56), aconitate (−0.36 to −0.90), and citrate/isocitrate (−0.59 to −1.36).SD alone showed mixed effects: depletion of G1P but modest increases in acetyl-CoA (+0.61 to +1.11), aconitate (−0.39 to +0.33), and citrate/isocitrate (+0.20 to +0.68) contents. The SDCQ combination produced the most pronounced suppression, with severe depletion of G1P, acetyl-CoA, aconitate, and citrate/isocitrate. B) SDPY: PY alone consistently reduced the levels of the glycolytic intermediates2,3-diphosphoglyceric acid (0.4 to −1.45-fold), malate (−0.28 to −0.56), and pyruvate derivatives (PD, −0.4 to −0.5). SD alone induced modest increases in malate and PD, with near-neutral effects on 2,3-diphosphoglyceric acid. The SDPY combination resulted in sustained depletion across all three metabolites (−0.23 to −0.83-fold for 2,3-diphosphoglyceric acid; −0.27 to −0.56-fold for malate; and −0.44--to −0.65-fold for PD).

**Figure 5.**
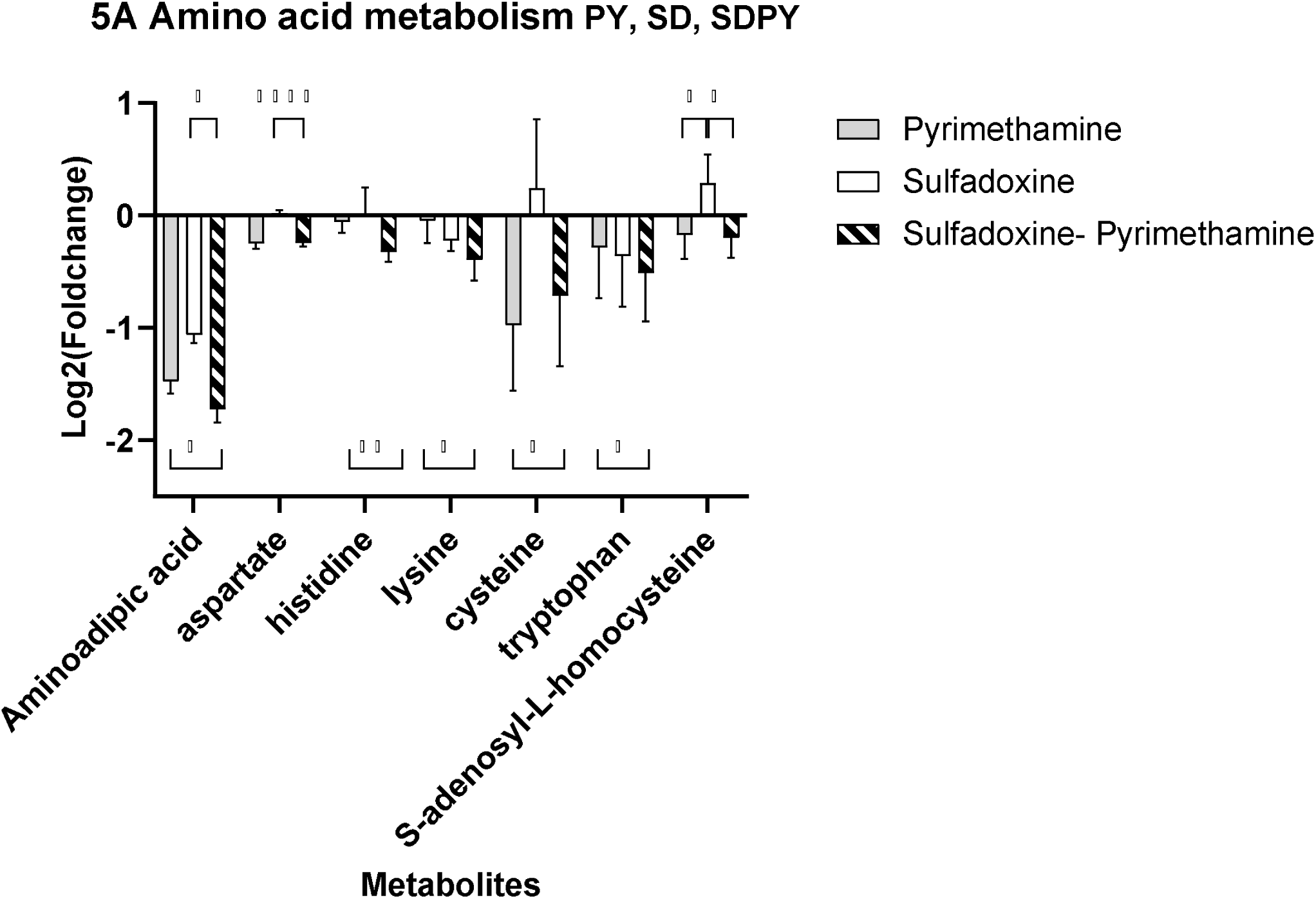
Amino acid metabolism in the case of isolated drugs and their combinations SDCQ and SDPY. A) SDPY: Aminoadipic acid was consistently downregulated across treatments, with the strongest reduction observed in the combined condition (−1.92 to −1.52). Aspartate showed minimal variation, fluctuating close to zero for SD. Histidine remained decrease under PY but increase under SD, and the combination induced a slight decrease. Lysine displayed mild negative shifts under PY and SD, with a greater reduction in the combined treatment (−0.75 to −0.14). Cysteine exhibited heterogeneity, with PY inducing both negative and positive changes, SD production increases in some replicates, and the combination trending toward decreases. Tryptophan showed mixed responses: PY reduced in most replicates, SD had variable effects, and the combination produced largely negative values. S-adenosyl-L-homocysteine exhibited variable regulation, increasing under SDs and decreasing under the combined treatment. In the SDCQ dataset, no significant differences were detected for the amino acid metabolites analyzed, suggesting a neutral effect of this combination on amino acid metabolism.

**Figure 6.**
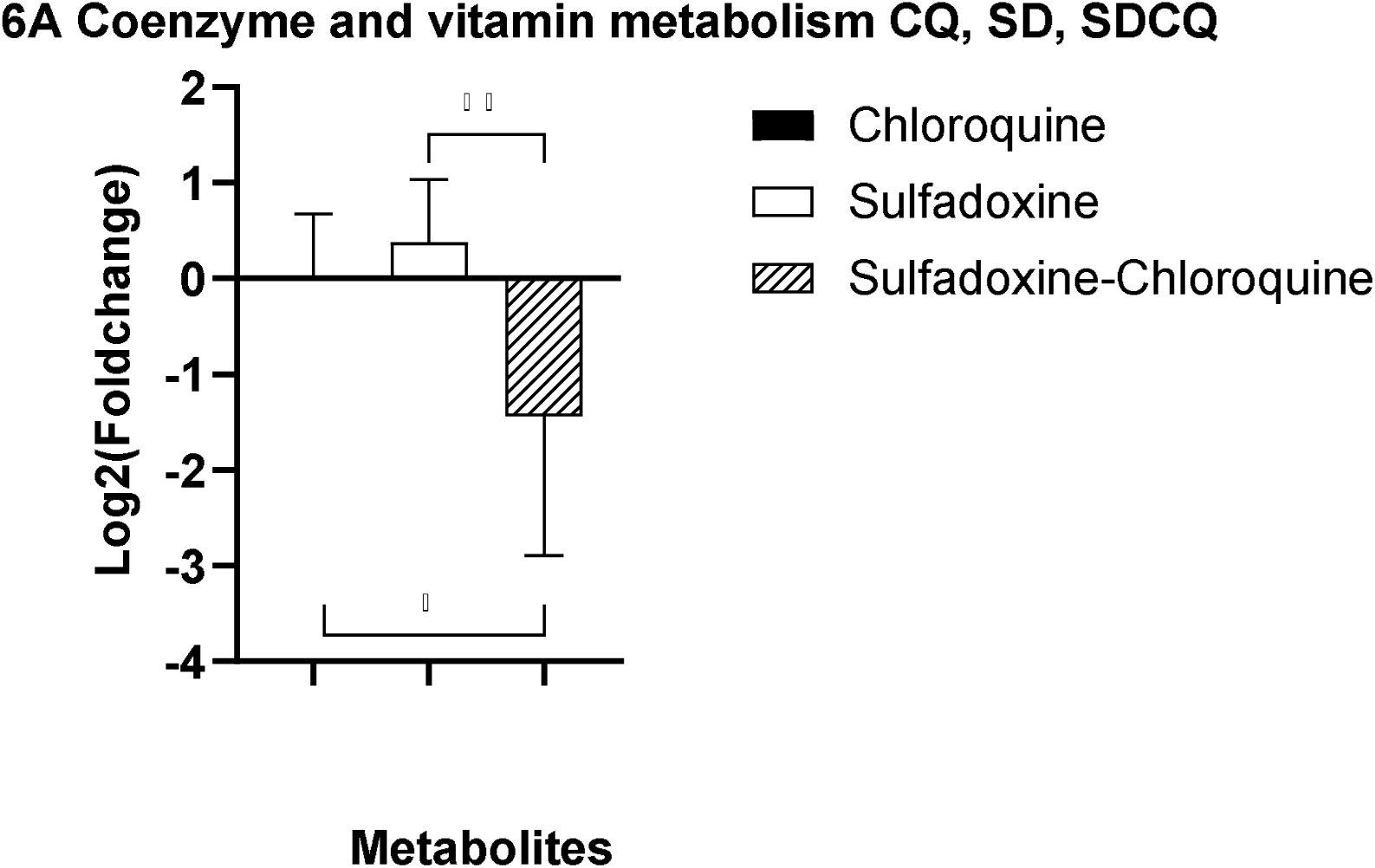
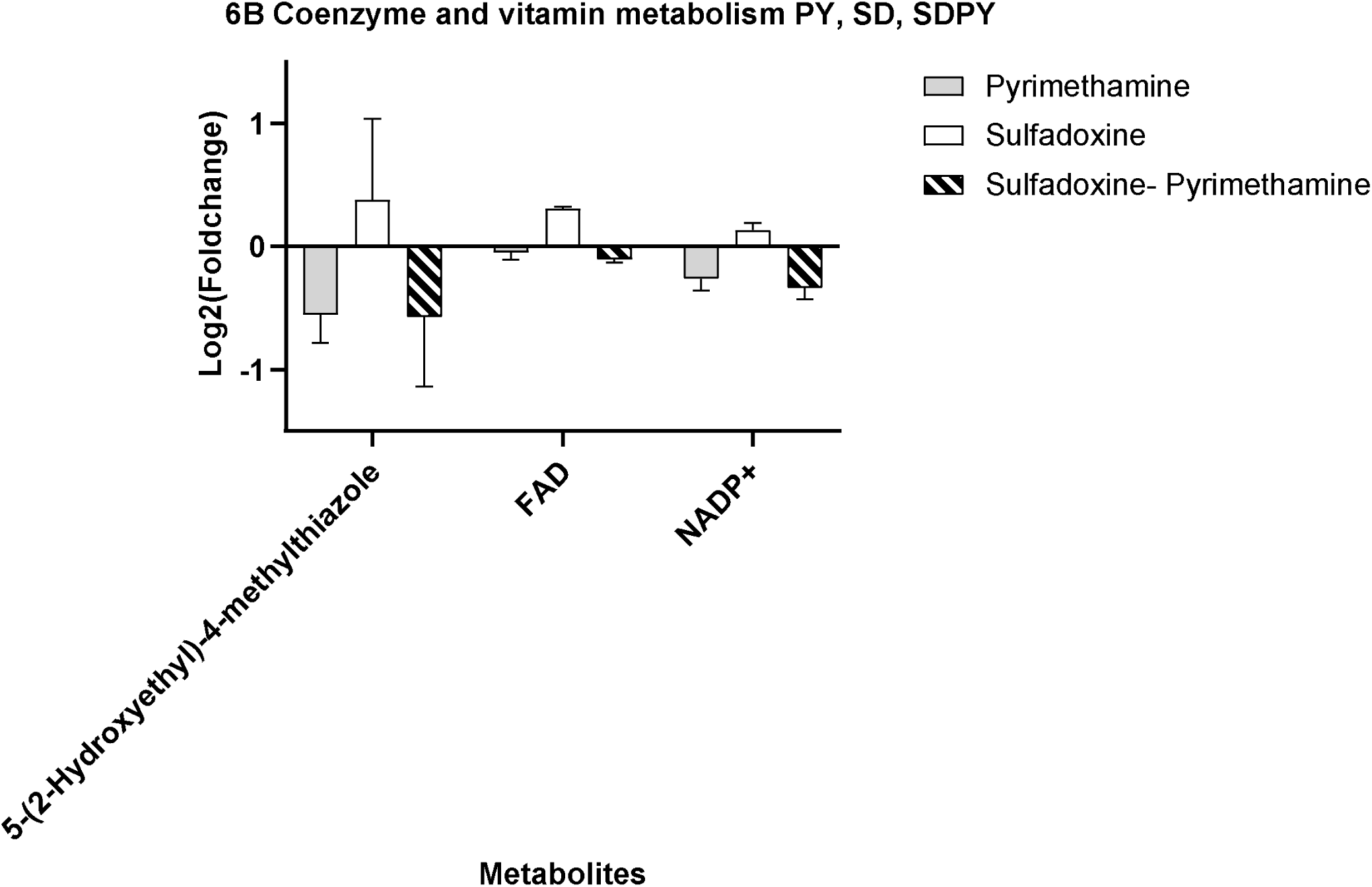
Coenzyme and vitamin metabolism in the case of isolated drugs and their combinations SDCQ and SDPY. A) SDCQ: For 5-(2-hydroxyethyl)-4-methylthiazole, CQ, SD, and their combination gave variable results inside tests; positive and negative. SDCQ markedly downregulated 5-(2-hydroxyethyl)-4-methylthiazole. B) SDPY: The thiazole derivative 5-(2-hydroxyethyl)-4-methylthiazole showed strong downregulation under PY and a variable response under SD treatment. The combination treatment resulted in suppression. FAD remained mostly stable under PY, increased moderately with increasing SD (0.27 to 0.33), and slightly decreased under the combination (−0.10 to −0.15). NADP+ was downregulated under PY (–0.36 to –0.06), upregulated under SD (0.04 to 0.24), and decreased under the combination treatment (–0.52 to –0.26).

**Figure 7.**
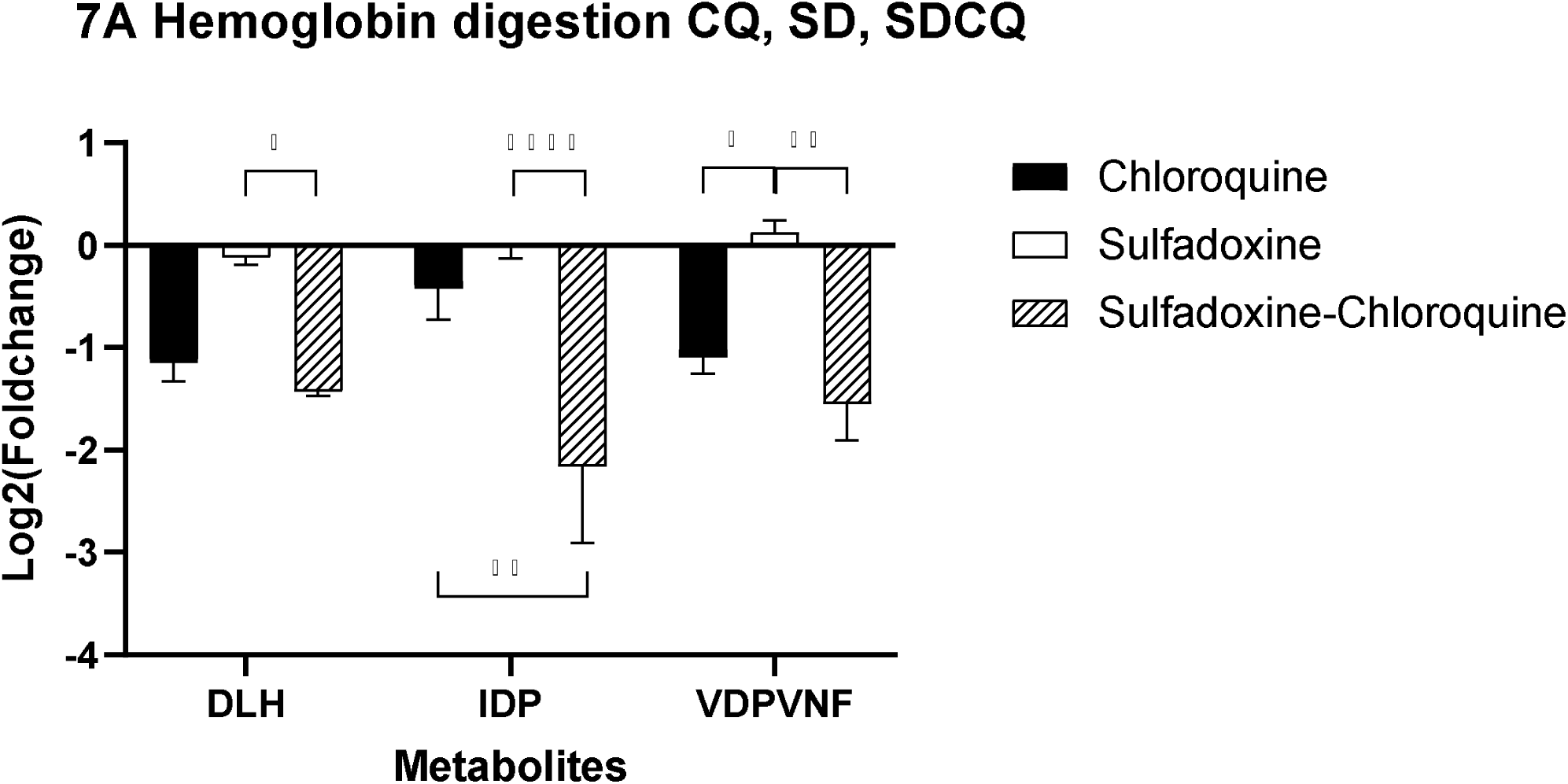
Effects on hemoglobin digestion in the case of isolated drugs and their combinations SDCQ and SDPY. A) SDCQ: CQ, SD, and their combination indicated strong reductions in DLH and IDP under chloroquine alone. SD alone produced mild effects, with slight increases in VDPVNF. However, the SDCQ combination resulted in marked decreases across DLH, IDP, and VDPVNF.

**Figure 8.**
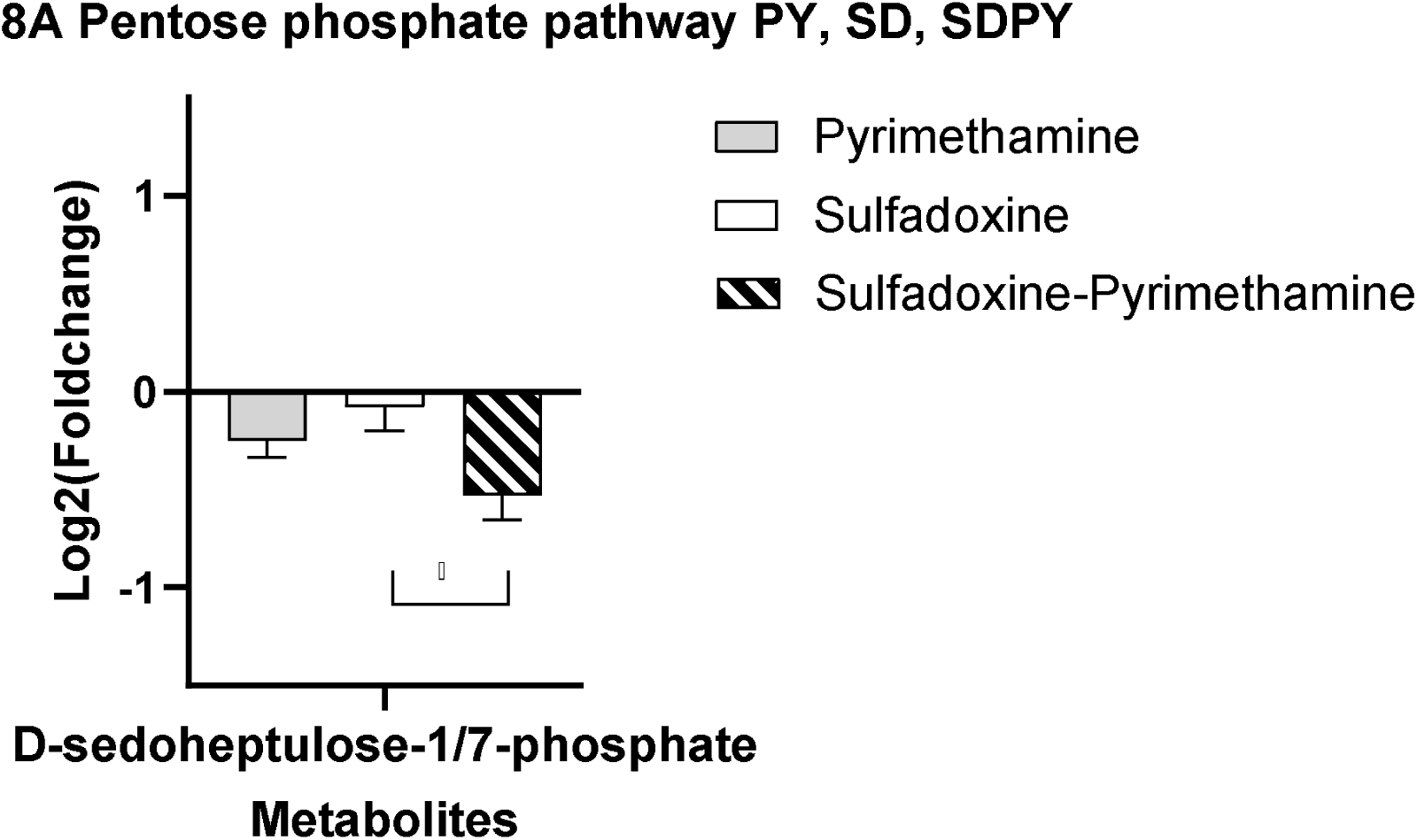
Pentose phosphate pathway metabolites in the case of isolated drugs and their combinations SDCQ and SDPY. A) SDPY: PY alone produced moderate decreases in D-sedoheptulose-1/7-phosphate (−0.408 to −0.114). SD alone had variable effects, slightly increasing or mildly decreasing the metabolite content (−0.319 to 0.056), indicating weaker activity. The combination of SDPY resulted in strong decreases across replicates, indicating a synergistic effect on reducing D-sedoheptulose-1/7-phosphate levels. SDCQ: No significant differences were observed, indicating limited or neutral metabolic effects when CQ and SD are combined for the pentose phosphate pathway.

**Figure 9.**
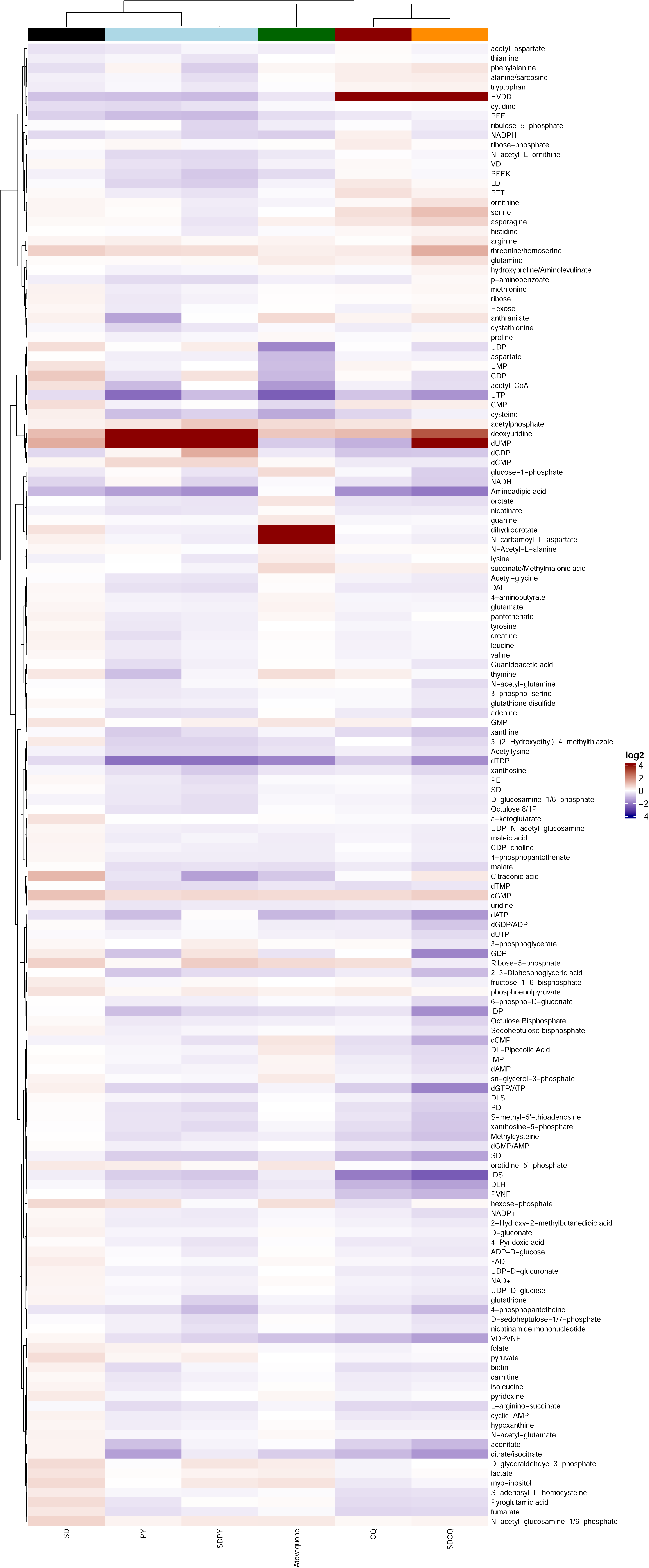
Metabolomic profiling of the compounds. Heatmap of the log2-fold change values of 154 metabolites, relative to the no-drug control. All the displayed log2-fold changes are the averages of three technical replicates for each compound tested. Distance clustering was performed using the Pearson**□**Ward method. The selected metabolites are denoted on the right, corresponding to specific metabolic signatures within a particular MoA class, and are color coded. Atovaquone, the hydroxynaphthoquinone component in Malarone®, was used as a control.

### Nucleotide and pyrimidine pathway perturbations

Nucleotide metabolism was markedly affected (Figure 2A, 2B), with single drug treatment showing distinct but complementary effects. For example, CQ induced strong dCDP depletion, while SD caused CMP and ADP-D-glucose accumulation. When combined (SDCQ), these effects were neutralized, highlighting potential metabolic buffering. PY alone dramatically increased the levels of deoxyuridine (10-fold) and dUMP (13-14-fold) (Figure 3A, 3B), which was consistent with the blockade of DHFR-dependent thymidylate synthesis. These metabolite signatures echo previous reports of antifolate-induced pyrimidine stress [32]. Importantly, a combination involving PY (SDPY) sustained or accentuated this accumulation, underscoring the robustness of the antifolate effect.

### Central carbon metabolism and the TCA cycle

CQ treatment alone broadly suppressed glycolytic and TCA intermediates(Figure 4A, 4B), including glucose-1-phosphate, citrate, and aconitate, suggesting global energy stress. SD, by contrast, displayed mixed effects, with modest increases in the levels of TCA intermediates. Drug combinations, particularly SDCQ, induced the most severe depletion across glycolytic and TCA metabolites, indicating additive inhibition of central carbon metabolism. Similar patterns of energy depletion under antifolate or quinoline pressure have been described in global metabolomics surveys (Allman *et al*., 2016), highlighting metabolic collapse as a shared mechanism of antimalarial toxicity [32].

### Amino acid metabolism

PY strongly reduced aminoadipic acid, acetylated amino acid derivatives, and cysteine, while SD produced more modest and variable effects (Figure 5A). Combination treatments generally accentuated these reductions, suggesting a compounded disruption of amino acid pools. These findings mirror broader metabolomic evidence that antifolate stress diverts amino acid metabolism toward compensatory one-carbon flux [35]. Interestingly, the SDCQ dataset showed no major amino acid alterations, indicating that this combination exerts its primary impact elsewhere.

### Coenzymes and vitamin metabolism

SD increased the contents of 5-(2-hydroxyethyl)-4-methylthiazole, FAD and NADP+ (Figure 6A, 6B). The combined treatment of SDCQ and SDPY tended toward depletion, reflecting impaired redox capacity. This finding aligns with prior findings that antifolate treatment perturbs cofactor balance and reduces antioxidant defenses [32, 41].

### Hemoglobindigestion

Peptides harvested from hemoglobin such as DLH, IDP, and VDPVNF were strongly reduced under CQ, and the effect was amplified by combinations with SD (Figure 7A). These observations support that CQ affects a known drug target, the digestive vacuole.

### Pentose phosphate pathway (PPP)

PY alone reduced D-sedoheptulose-1/7-phosphate and, when combined with SD, produced a synergistic decrease (Figure 8A). The depletion of sedoheptulose-1/7-phosphate suggests reduced NADPH production and ribose-5-phosphate availability, thereby limiting nucleotide biosynthesis and antioxidant defense. These findings parallel recent metabolomics reports that highlight PPP inhibition as a vulnerability under antifolate and oxidative stress conditions [32, 35, 41].

### Metabolic fingerprint profiling

To gain further insight into the biochemical pathways affected by the drug combinations, metaprint pathway analysis was performed on the 154 targeted metabolites measured by LC**□**MS (Figure 9). Metabolic fingerprint profiling revealed that combination therapies produced broader and more synergistic signatures than single drugs did, reflecting amplified metabolic pressure (Figure 10). The suprahexagonal metaprint highlights overlapping yet distinct metabolic modules compared with atovaquone controls, confirming unique drug-specific metabolic fingerprints [32].

**Figure 10.**
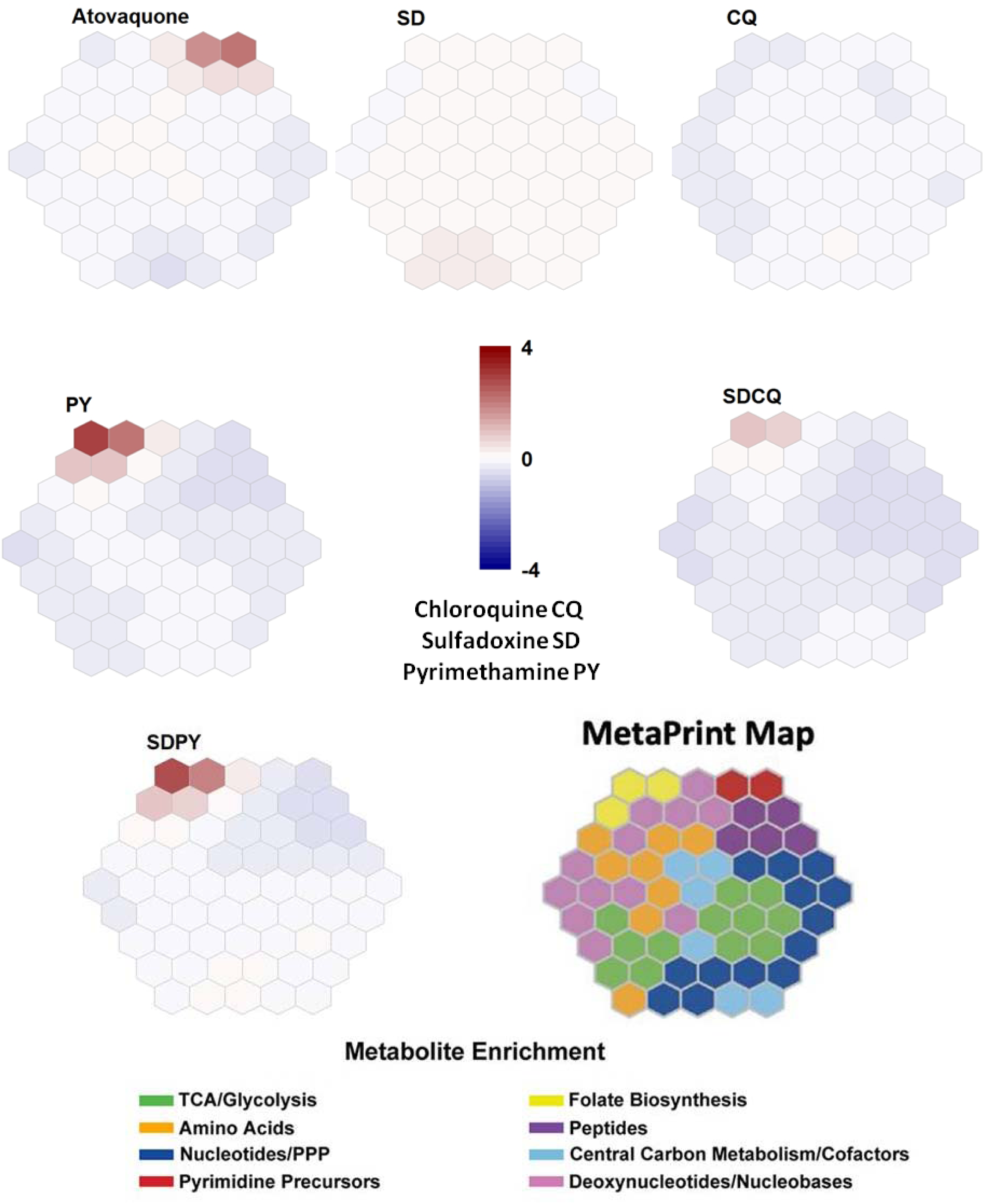
Metabolomic analysis visualized through a suprahexagonal fingerprint. The metabolites were grouped into eight broad KEGG-based pathways and color-coded on the base map. The metabolic signature of atovaquone is shown as a reference, highlighting key distinguishing metabolites. (See the metabolite mapping and the metaprint map with enrichment from Allman *et al*., 2016) [32].

## Discussion

This study provides an assessment of the effects of pairing antimalarial drugs against *Plasmodium falciparum* by comparing treatment with CQ, PY, SD, and their combinations SDCQ and SDPY. Dose□response analysis confirmed the high potency of PY and CQ against *P. falciparum*, both with nanomolar IC₅₀ values (Figure 1). These findings are consistent with earlier studies showing that antifolates and quinoline derivatives achieve strong growth suppression at low nanomolar concentrations [42, 43, 44, 45]. In contrast, sulfadoxine required micromolar concentrations to inhibit parasite growth, highlighting its relatively weaker standalone efficacy, which is in line with its role as a synergistic partner in combination therapies [46]. This difference highlights mechanistic differences: PY interferes with the function of human DHFR, an enzyme essential for folate metabolism. Since DHFR drives the synthesis of purines and thymidine, the inhibition of DHFR by antifolates such as methotrexate leads to a reduction in the cell’s supply of dTTP, ATP, and GTP [47], while CQ exerts its antimalarial action by blocking the detoxification of free heme released during hemoglobin digestion, preventing its conversion into inert hemozoin. The accumulation of toxic heme within the digestive vacuole leads to parasite death [48]. Sulfadoxine, a sulfa-based antifolate, blocks dihydropteroate synthase (DHPS) by mimicking para-aminobenzoic acid (pABA).This disruption halts folate production, impairing nucleotide synthesis and amino acid metabolism and ultimately leading to parasite death [49]. The pharmacological rationale for therapies such as SDPY or SDCQ is reinforced by evidence that the high nanomolar potency of PY and CQ compensates for the micromolar activity of SD, resulting in multiple points of attack that enhance antiplasmodium efficacy.

By using targeted mass spectrometry to measure a wide range of metabolites from central metabolism, nucleotide biosynthesis, amino acid turnover, cofactors and redox balance, the pentose phosphate pathway (PPP), and glycan/lipid precursors, both drug-specific and synergistic metabolic effects were uncovered that explain the superior efficacy of combinations.

Single drug treatments produced distinct perturbations in nucleotide and folate metabolism (Figures 2,3). PY treatment led to the accumulation of deoxyuridine and dUMP due to the inhibition of dihydrofolate reductase (DHFR), an enzyme essential for regenerating tetrahydrofolate (THF) required in one-carbon transfer reactions. This disruption blocks the conversion of dUMP to dTMP, impairs thymidylate synthase activity, and interrupts folate-dependent pyrimidine biosynthesis, ultimately halting DNA synthesis and suppressing *Plasmodium falciparum* replication [44]. Allman and coworkers stressed that DHFR-TS inhibitors showed increased dUMP and NADPH, which are unique signatures and cluster compounds P218 and MMV667487 [32]. SD treatment led to perturbations in pyrimidine intermediates, including mild accumulation of dUMP and increased N-carbamoyl-L-aspartate levels. CQ induces only modest perturbations in pyrimidine nucleotide pools; its metabolic signature, marked by DNA intercalation and oxidative stress, is distinct from that of DHFR/DHPS inhibitors and far less disruptive to pyrimidine biosynthesis than antifolates [32].

With respect to paired drugs, SDCQ treatment strongly increased the level of deoxyuridine monophosphate (dUMP), with downstream perturbations suggesting reinforced DHFR-TS inhibition, a known hallmark of antifolate action [22, 32]. A similar disruption of dual therapy has been observed with chalcone and artemisinin in chloroquine-sensitive *Plasmodium falciparum* strains [50]. The increase in dUMP, which is converted to deoxythymidine monophosphate (dTMP), suggested an enhancement of DHFR-TS production. dUMP upregulation has been shown to lead to blood and liver tissue thymidylate synthase (TS) inhibition in patients with primary tumors and liver metastases [51]. Drugs such as the antibiotics trimethoprim, WR99210, and methotrexate, structural analogs of folate, are known to possess antimalarial activity through this mode of action (MoA) [32, 52, 53]. SDCQ prevented SD-induced CMP and ADP-D-glucose accumulation, indicating flux neutralization and redistribution of carbon precursors away from pyrimidine synthesis.

Considering central carbon metabolism and the TCA cycle (Figure 4), PY consistently suppressed glycolytic intermediates (2,3-DPG, PD) and TCA metabolites (malate) secondary to folate-dependent NADPH depletion [32, 44] (Figure 3). SD treatment resulted in partial compensation of TCA flux by increasing acetyl-CoA, aconitate, and citrate levels, despite G1P depletion, suggesting impaired glycolytic entry. CQ depleted TCA cycle intermediates, corroborating the findings of Gao and colleagues that CQ disrupts parasite glycolysis and energy metabolism to exert its antimalarial effects [54]. SDCQ induced dramatic collapse of both glycolysis and TCA intermediates (acetyl-CoA, aconitate, citrate/isocitrate), reflecting synergistic antifolate-CQ disruption of energy metabolism [55]. In *P. falciparum*, citrate and isocitrate are essential TCA cycle components that play roles beyond energy production, including responses to oxidative stress [56]. The TCA cycle begins with the formation of citrate from acetyl-CoA and oxaloacetate, which is catalyzed by citrate synthase. Artemisinin-resistant parasites accumulate less citrate in the presence of drugs [57]. Aconitase catalyzes the reversible conversion of citrate to isocitrate via aconitate. Aconitase inhibition can disrupt the TCA cycle, hindering parasite energy production [58]. SDPY suppressed parasites’ ability to break down glucose for energy and downstream respiration, which is consistent with the dual blockade of folate-mediated carbon metabolism [59].

PY and SD, individually and in combination, consistently reduced amino acid metabolites, including lysine, cysteine, tryptophan, and aminoadipic acid, indicating perturbation of catabolic and biosynthetic pathways (Figure 5). Murithi and collaborators reported that CQ induced a >2-fold increase in N-acetyl-lysine, which was not observed under piperaquine pressure [45]. SDCQ showed neutralizing trends, suggesting complex compensatory interactions.

Considering cofactors and redox metabolism (Figure 6), PY suppressed thiazole derivatives, NADP+, 4-phosphopantetheine, thiamine, and glutathione, reflecting a disruption in folate-dependent redox. SD treatment often resulted in the upregulation of FAD, NADP+, and thiazole derivatives, indicating stress-induced compensatory responses. Thiamine pyrophosphate (TPP), the active form of vitamin B1, is a crucial cofactor for pyruvate dehydrogenase (PDH), which links glycolysis and the tricarboxylic acid (TCA) cycle. TPP synthesis involves the merging of pyrimidine and thiazole branches. The thiazole branch involves the phosphorylation of 5-(2-hydroxyethyl)-4-methylthiazole (THZ) to THZ phosphate (THZ-P) by THZ kinase (ThiM). *Plasmodium falciparum* synthesizes THZ during thiamine biosynthesis [60]. THZ and acetyl-CoA are under-regulated with SDCQ compared to SD, and THZ comparared to CQ, potentially impacting parasite vitamin B1 production.

The metabolic signature of decreased DLH peptide levels DLH, IDP, and HVDD is among those related to the inhibition of hemoglobin endocytosis and/or catabolism within the digestive vacuole, as previously observed across multiple antimalarial compounds (Figure 7) [45]. A reduction in peptide substrates such as HVDD and VDPVNF indicates disruption of hemoglobin catabolism and amino acid supply for anabolic processes, which is consistent with PfA-M1- and PfCRT-mediated transport inhibition [61, 62]. The peptide VDPVNF was previously identified asa putative PfCRT substrate in [^3^H]-CQ *cis*-inhibition experiments in the *Xenopus laevis* oocyte model system and was downregulated in SDCQ compared with SD. VDPVNF(VL-6, a hemoglobin fragment), which can be degraded by the *P. falciparum* M1 alanyl-aminopeptidase (*Pf*A-M1) (an enzyme involved in the terminal stages of hemoglobin digestion and the generation of an amino acid pool within the parasite), was also observed to inhibit CQ transport through *Pf*CRT [61, 62, 63].

PY treatment alone or in combination with SD reduced key pentose phosphate pathway (PPP) intermediate (D-sedoheptulose-1/7-phosphate) levels, limiting NADPH production and ribose-5-phosphate availability, which are essential for nucleotide and redox metabolism (Figure 8). SD had weak effects; SDPY amplified PPP suppression. Trophozoite surface glycoproteins possess truncated O- and N-glycans with 1-2 GlcNAc moieties of unknown function. N-glycans on the surface of infected RBCs decrease, and sugar nucleotide levels rise, suggesting that *Plasmodium* parasites increasingly utilize scavenging pathways to build up the required glycosylation [63]. The common precursors for glycan biosynthesis are sugar nucleotides, from which monosaccharides are transferred to nascent glycan chains, proteins or lipids by glycosyltransferases [65]. Sugar nucleotides generally consist of a nucleotide, such as UDP, GDP, CMP, or CDP, connected to a monosaccharide [66, 67]. The sugar nucleotides CDP, UDP, and GDP were downregulated under SDCQ relative to SD, confirming that the inhibition of glycan synthesis via impaired glycosyltransferase activity affects parasite survival. This reduction confirms that the combined treatment acts on glycan synthesis by reducing the levels of sugar nucleotides that can be transferred by glycotransferases. In eukaryotes, UDP-glucose (UDP-Glc) is synthesized from glucose-6-phosphate (Glc6P) and glucose-1-phosphate (Glc1P). Glc1P is then activated to UDP-Glc, typically by UTP-glucose-1-phosphate uridylyltransferase [68]. The pronounced decrease in nucleoside di- and tri-phosphate levels described (UDP, GDP, dGTP/ATP) for SDCQ has been identified as a signature of Na+/H+-dependent ATPase PfATP4 inhibition, as described previously for compounds KAE609 and MMV019017 [32].

Four distinct metabolic "signatures" to drug class exposure were previously reported by Allman *et al*. (2016), each characterized by disruptions in one of the following pathways: folate biosynthesis, cellular homeostasis, mitochondrial electron transport chain and pyrimidine synthesis, or hemoglobin degradation [32]. These findings indicate that compound mixtures that act on distinct pathways continue to exhibit unique metabolic signatures, although with modified intensities [32]. The metabolic fingerprint analysis highlights distinct but converging effects of the tested antimalarial agents and their combinations (Figure 10). PY produces a pronounced signature centered on pyrimidine precursors, which is consistent with its role as a dihydrofolate reductase inhibitor that blocks folate-mediated nucleotide biosynthesis [32]. In contrast, SD alone elicited a weaker perturbation, reflecting its comparatively modest impact when dihydropteroate synthase was inhibited in isolation. CQ showed minimal alterations across central metabolic pathways, in agreement with its primary mechanism of disrupting heme detoxification rather than carbon or nucleotide flux [32]. The combination of SD and PY (SDPY) produced the most consistent antifolate fingerprint, characterized by strong suppression of pyrimidine precursor pathways. This outcome aligns with the well-established dual blockade of folate metabolism by targeting both DHPS and DHFR-TS [52]. In contrast, the SDCQ pairing showed only modest and variable changes, which mirrors clinical findings that this combination reduces consistent efficacy. This observation is corroborated by van den Broek and collaborators in comparison with artemisinin-based combination treatments such as mefloquine + artesunate or lumefantrine + artemether [69].

Taken together, these results indicate that antifolate-driven suppression of pyrimidine metabolism remains the dominant metabolic outcome in combination therapies, with CQ acting mainly as a stress enhancer rather than a direct metabolic disruptor. These fingerprints reinforce the value of targeting pyrimidine biosynthesis as a metabolic bottleneck in malaria parasites and help explain why antifolate-based combinations remain highly effective despite varying levels of resistance.

These findings demonstrate that combination therapies act on complementary metabolic nodes, producing distinct and amplified metabolic stress signatures. Compared with monotherapies, SDPY and SDCQ combinations achieve greater inhibition of nucleotide, carbon, amino acid, cofactor, PPP, and glycan metabolism. This provides mechanistic justification for dual regimens, supporting enhanced efficacy and potential mitigation of resistance [70, 71, 72].

## Conclusions

This study demonstrates that the integration of metabolomics with pharmacological profiling provides a powerful framework for understanding how antimalarial drug combinations disrupt *Plasmodium falciparum* metabolism. Across all treatments, combination therapies produced broader and more intense metabolic disturbances than single agents did, confirming that synergistic drug interactions operate through coordinated, pathway-specific perturbations. These effects mechanistically explain their superior antimalarial potency relative to that of monotherapies.While sulfadoxine alone induced compensatory increases in several redox cofactors, its combination with chloroquine or pyrimethamine produced variable outcomes, reflecting complex redox reprogramming. A central finding is the strong and recurrent inhibition of the pentose phosphate pathway, especially with pyrimethamine-containing combinations. This dual suppression of folate metabolism and PPP flux appears critical for limiting nucleotide availability and reducing the parasite’s capacity to counter oxidative stress. Moreover, pyrimidine pathway disruption has emerged as the hallmark metabolic signature of effective antifolate-based therapy.Together, our results provide mechanistic evidence supporting the clinical advantages of rational antimalarial combinations.

## List of Abbreviations

ACT: Artemisinin-based Combination Therapy
ADP: Adenosine Diphosphate
ATP: Adenosine Triphosphate
CMP: Cytidine Monophosphate
CDP: Cytidine Diphosphate
CQ: Chloroquine
dCDP: Deoxycytidine Diphosphate
dGTP: Deoxyguanosine Triphosphate
DHFR: Dihydrofolate Reductase
DHFR-TS: Dihydrofolate Reductase–Thymidylate Synthase
DHPS: Dihydropteroate Synthase
DMSO: Dimethyl Sulfoxide
dTMP: Deoxythymidine Monophosphate
dUMP: Deoxyuridine Monophosphate
dTTP: Deoxythymidine Triphosphate
DV: Digestive Vacuole
FAD: Flavin Adenine Dinucleotide
FPIX: Ferriprotoporphyrin IX
G1P: Glucose-1-Phosphate
Glc6P: Glucose-6-Phosphate
GDP: Guanosine Diphosphate
GTP: Guanosine Triphosphate
HPLC: High-Performance Liquid Chromatography
IRS: Indoor Residual Spraying
ITN: Insecticide-Treated Net
LC-MS: Liquid Chromatography-Mass Spectrometry
MPAG: Malaria Policy Advisory Group
NADP⁺: Nicotinamide Adenine Dinucleotide Phosphate ion
NADPH: Nicotinamide Adenine Dinucleotide Phosphate
PABA: para-Aminobenzoic Acid
PBS: Phosphate-Buffered Saline
PfCRT: Plasmodium falciparum Chloroquine Resistance Transporter
PfATP4: Plasmodium falciparum Na⁺/H⁺-Dependent ATPase
PfA-M1: Plasmodium falciparum Alanyl-Aminopeptidase M1
PPP: Pentose Phosphate Pathway
PY: Pyrimethamine
RDT: Rapid Diagnostic Tests
RPMI: Roswell Park Memorial Institute Medium
SAGE: Strategic Advisory Group of Experts
SD: Sulfadoxine
SDCQ: Combination of Sulfadoxine + Chloroquine
SDPY: Combination of Sulfadoxine + Pyrimethamine
SEM: Standard Error of the Mean
SMC: Seasonal Malaria Chemoprevention
TCA: Tricarboxylic Acid Cycle
THF: Tetrahydrofolate
TPP: Thiamine Pyrophosphate
UDP: Uridine Diphosphate
UDP-Glc: Uridine Diphosphate-Glucose
UTP: Uridine Triphosphate

## Acknowledgments

EMF gratefully acknowledges the Fulbright Program for awarding a prestigious Fulbright Academic Fellowship. The Llinás lab is grateful for support from The Pennsylvania State University and the Huck Institutes for the Life Sciences. We also acknowledge the Huck Institutes’ Metabolomics Core Facility (RRID:SCR_023864) for maintenance of the ThermoExactive Plus used in this study.

## Author

EMF, TQ, JM, and ML. designed,performed experiments and analyzed data. EMF, TQ, JM, and ML. wrote the manuscript and generated figures. ML provided critical comments, project and funding management. All authors wrote, read, and approved the final manuscript.

## Data availability

Data available in the NIH data repository, Metabolomics Workbench Study project doiPR002764, and Study ID ST004358, available at https://dev.metabolomicsworkbench.org:22222/data/DRCCMetadata.php?Mode=Study&StudyID=ST004358&Access=WubV3794

## Declarations

### Ethics approval and consent to participate

Not applicable

### Consent for publication

Not applicable.

